# The paradoxical impact of drought on West Nile virus risk: insights from long-term ecological data

**DOI:** 10.1101/2025.01.21.634155

**Authors:** Samantha Sambado, Terrell J Sipin, Zoe Rennie, Ashley Larsen, James Cunningham, Amy Quandt, Dan Sousa, Andrew J MacDonald

## Abstract

Mosquito-borne diseases are deeply embedded within ecological communities, with environmental changes – particularly climate change – shaping their dynamics. Increasingly intesense droughts across the globe have profound implications for the transmission of these diseases, as drought conditions can alter mosquito breeding habitats, host-seeking behaviors, and mosquito-host contact rates. To quantify the effect of drought on disease transmission, we use West Nile virus (WNV) as a model system and leverage a robust mosquito and virus dataset consisting of over 500,000 trap nights collected from 2010-2023, spanning a historic drought period followed by atmospheric rivers. We pair this surveillance dataset with a novel modeling approach that incorporates monthly changes in bird host community competence, along with drought conditions, to estimate the effect of drought severity on WNV risk using panel regression models. Our results show that while drought decreases mosquito abundances, it paradoxically increases WNV infection rates. This counterintuitive pattern likely stems from reduced water availability, which concentrates mosquitos and pathogen-amplifying bird hosts around limited water sources, thereby increasing disease transmission risk. However, the magnitude of the effect depends critically on mosquito species, suggesting species-specific behavioral traits are key to understanding the effect of drought on mosquito-borne disease risk across real landscapes.

## BACKGROUND

Climate change and extreme climate events are a major threat to human health (1,2). Human health risks, such as vector-borne diseases, are particularly sensitive to changes in climate and are an increasing health threat in North America (3,4). Ectothermic vectors that rely on water for breeding, such as mosquitos, are sensitive to changes in temperature and the availability of standing water (5,6). Drought, drier-than-normal conditions, can reduce the availability of standing water, thereby potentially affecting mosquito development rates and breeding success (5,7). Many diseases spread by mosquitos involve various species of vectors and hosts, which may respond differently to drought even when they coexist within the same community (8,9). To identify the effects of drought on mosquito-borne disease risk, it is essential to differentiate between species-specific responses, enabling more nuanced vector control strategies and public health messaging.

Drought is an important driver of ecosystem dynamics (10,11) which affects mosquito populations that rely on water to complete their life cycle (7,12,13). Reduced precipitation can lead to diminished habitats for egg laying, thereby decreasing mosquito abundances (5,7,12). However, reduced precipitation can decrease the total area of standing water, potentially concentrating mosquitos and hosts that can become infected and transmit infections back to mosquitos (7,12). Increased contact rates between mosquitos and important hosts of pathogens can increase pathogen transmission rates (7,14). In drought-affected regions, natural waterways like rivers can provide essential water resources across the landscape (8). However, rivers that rely on snowmelt can experience substantial fluctuations depending on the timing and rate of snowmelt, which are influenced by drought conditions (12,14–16). Parsing apart the potentially conflicting impacts of drought on two critical metrics for mosquito-borne disease risk – mosquito abundance and infection rates – will be essential as droughts are projected to become more frequent and severe (7,11).

West Nile virus is the most common mosquito-borne disease in North America since its introduction to the United States in 1999 (4,17). Given that West Nile virus relies on multiple factors such as mosquito abundances, the presence of disease-carrying hosts such as birds, and human contact rates, a mechanistic understanding of drought effects is crucial for predicting mosquito-borne disease accurately (18). West Nile virus is transmitted by many *Culex* spp. mosquitos and primarily maintained by bird hosts (19). Certain bird species, known as “competent hosts”, can acquire and maintain the virus, serving as infectious blood meals for mosquitos. In humans, infections are mostly asymptomatic (∼80% of known cases) but they can be severe, resulting in neuroinvasive disease in about ∼1% of known cases (4). Over the past twenty years, California has accounted for 15% of all reported human cases of West Nile virus in the United States, with the Central Valley reporting incidence rates that are roughly twice the state average (4,17,20). This is thought to be due to extensive agricultural production requiring irrigation, which enhances mosquito breeding habitat (6). Agricultural regions may also have higher concentrations of lower income and disadvantaged communities that face challenges such as substandard housing, inequitable access to healthcare, and inadequate water management, all of which can increase human health risks (20–23).

Situated in the southern part of California’s Central Valley, Kern County has, on average, less than 153 mm of precipitation per year and has experienced multiple historic droughts over the past two decades (10,24). During that time, Kern County has also experienced persistent and in some years, relatively high rates of West Nile virus activity. The county’s landscape is predominantly agricultural, centered around the major city of Bakersfield with a population of approximately 400,000 (United States Census Bureau 2024). Of the 2.3 million acres of classified farmland, roughly 30% are irrigated (USDA National Agricultural Statistics Service). With contrasting rural agricultural and urban settings, there are two significant vectors of West Nile virus in these landscapes: *Culex tarsalis* and *Culex quinquefasciatus*, considered the primary rural and urban vector of WNV, respectively (14). The rural vector prefers breeding in larger bodies of water like seasonal wetlands, and experiences higher population numbers following an increase of water on the landscape, whereas the urban mosquito is more likely to breed in containers and small artificial water sources, as well as be flushed out by large pulses of water (8,12). The disease potential of these vectors also differs, where the rural mosquito can transmit higher pathogen loads but typically comes in less contact with humans compared to the urban mosquito (12). However, rural mosquitos are more likely to encounter vulnerable populations such as agricultural workers who may not have access to public health messaging in their native language or access to health care (21,22). Our study area, which has experienced recent and extreme variations in drought conditions, along with consistent, longitudinal mosquito surveillance activities, presents a compelling opportunity to explore the effects of drought on mosquito-borne disease risk.

In this study, we investigate the effects of drought severity on mosquito abundance and West Nile virus infection rates in Kern County, California between 2010-2023. We aim to build upon prior small-scale empirical studies that suggest drought differentially impacts two important mosquito-borne disease risk metrics – mosquito abundances and West Nile virus infection rates (12,14,25) – here using a novel data-driven approach. We use drought data and field surveillance data from 278 trap stations that were deployed over 500,000 trap nights resulting in 3.6 million mosquitos to estimate changes in mosquito abundance and WNV infection rates through time and across variable drought conditions to address the following questions:

1. How do *Cx. tarsalis* and *Cx. quinquefasciatus* abundances respond to drought severity?
2. How does WNV risk in *Cx. tarsalis* and *Cx. quinquefasciatus* respond to drought severity?

## METHODS

### Mosquito and WNV surveillance data

The Kern Mosquito and Vector Control District conducts regular mosquito surveillance with standardized trapping throughout the year, increasing their efforts during the summer months when mosquito activity peaks (8). The number of mosquitos is reported per trap night and include information on mosquito species, life stage, and sex. Mosquitos from the same trap, trap night and species are aggregated into pools and screened for WNV infection. For this study, we use data from the months of April through October during 2010-2023 and only on *Cx. tarsalis* and *Cx. quinquefasciatus* adult female mosquitos, the primary vectors of WNV in Kern County, and their respective WNV infection rates (14) (Table 1, Fig. 1).

**Figure 1.**
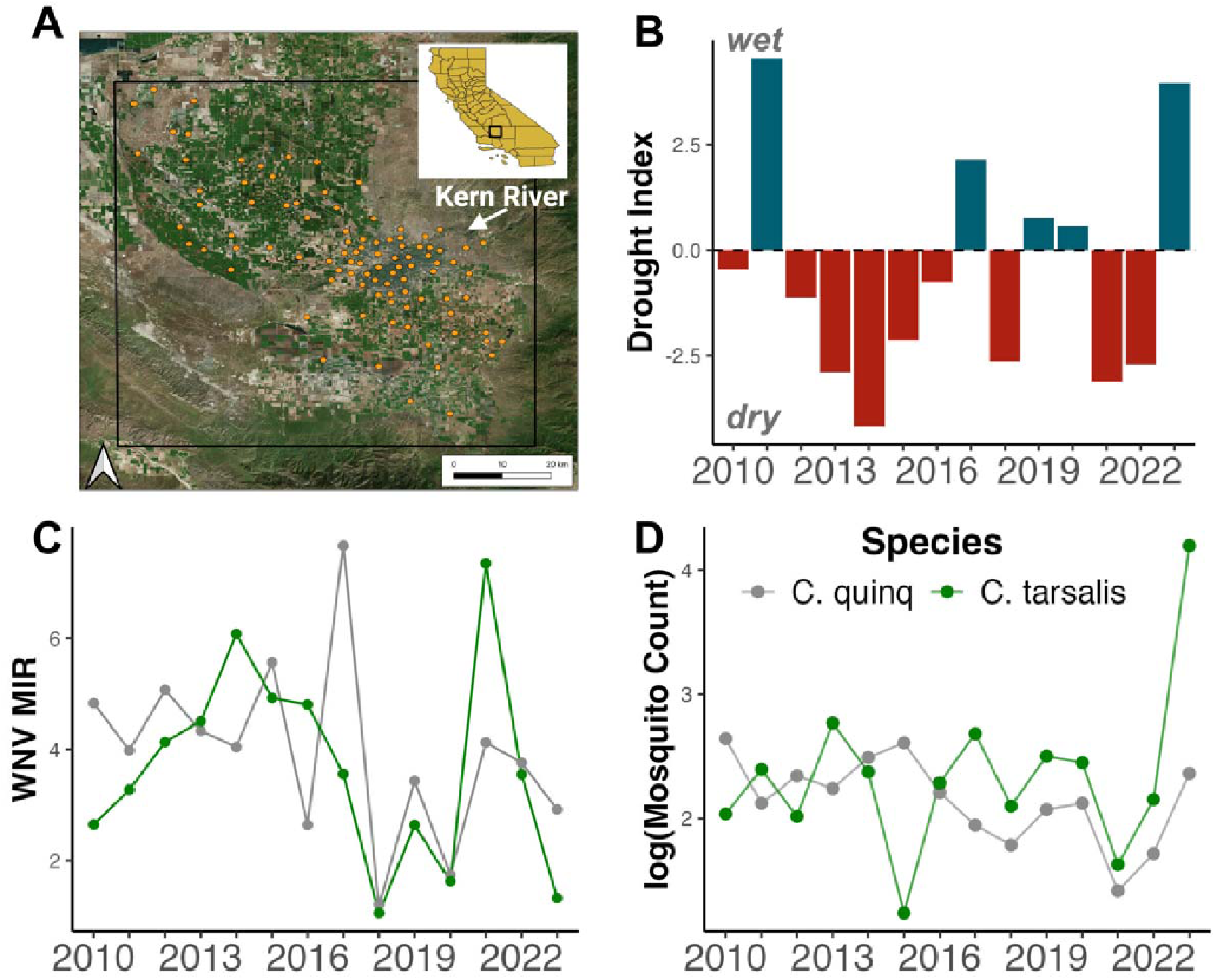
Study system: (A) clusterID locations centered around Bakersfield, CA, (B) yearly mean PDSI drought index, (C) yearly mean WNV minimum infection rate (MIR), and (D) log of yearly mean mosquito count per clusterID. In plots (C) and (D), mosquito species are color-coded: gray for *Cx. quinquefasciatus* (urban vector) and green for *Cx. tarsalis* (rural vector).

**Table 1.** Summary of yearly environmental characteristics and mosquito-borne disease metrics. Environmental characteristics include cumulative precipitation (Tot. Precip.), number of days that exceeded 30 °C (> 30), number of days that reach below 14 °C (< 14) within a calendar year. Mosquito-borne disease metrics are represented by yearly mean mosquito count and WNV minimum infection rate (MIR) per trap night with standard deviations per *Culex* species.

| Year | Environment | | | Mean count ± sd | | Mean WNV MIR ± sd | |
| --- | --- | --- | --- | --- | --- | --- | --- |
|  | Tot. Precip. | > 30 | < 14 | tarsalis | quinquefasciatus | tarsalis | quinquefasciatus |
| 2010 | 153 | 122 | 225 | 8.2 ± 39 | 15.1 ± 51 | 2.7 ± 8 | 4.8 ± 11 |
| 2011 | 259 | 116 | 239 | 11.4 ± 32 | 8.7 ± 19 | 3.3 ± 11 | 4.0 ± 12 |
| 2012 | 129 | 154 | 224 | 7.7 ± 47 | 10.6 ± 31 | 4.2 ± 11 | 5.1 ± 11 |
| 2013 | 82 | 142 | 236 | 16.5 ± 130 | 9.7 ± 25 | 4.4 ± 9 | 4.3 ± 10 |
| 2014 | 84 | 159 | 198 | 11.2 ± 58 | 12.5 ± 40 | 6.0 ± 13 | 4.0 ± 11 |
| 2015 | 128 | 150 | 191 | 3.5 ± 12 | 14.2 ± 80 | 4.6 ± 9 | 5.6 ± 11 |
| 2016 | 163 | 140 | 218 | 9.9 ± 79 | 9.2 ± 35 | 5.1 ± 11 | 2.6 ± 8 |
| 2017 | 221 | 136 | 227 | 14.8 ± 64 | 7.5 ± 18 | 3.4 ± 9 | 7.7 ± 15 |
| 2018 | 108 | 129 | 214 | 8.4 ± 62 | 6.4 ± 23 | 1.1 ± 5 | 1.2 ± 5 |
| 2019 | 192 | 129 | 225 | 12.8 ± 55 | 8.4 ± 25 | 2.4 ± 8 | 3.4 ± 8 |
| 2020 | 192 | 146 | 204 | 12.5 ± 108 | 9.3 ± 27 | 1.7 ± 6 | 1.7 ± 6 |
| 2021 | 82 | 137 | 222 | 5.6 ± 22 | 4.6 ± 17 | 7.3 ± 13 | 4.1 ± 11 |
| 2022 | 133 | 141 | 202 | 9.1 ± 71 | 6.0 ± 17 | 3.5 ± 10 | 3.8 ± 9 |
| 2023 | 320 | 112 | 224 | 69.4 ± 184 | 11.1 ± 30 | 1.1 ± 6 | 2.9 ± 10 |

To produce more consistent local estimates of abundance and WNV infection through time at an individual station, as well as account for spatial autocorrelation and non-independence of mosquito abundances in highly spatially aggregated trap stations, we spatially clustered nearby trap stations following the approach of MacDonald et al. 2024 (26). We cluster trap stations within ∼ 1,500 m radius based on the daily dispersal rate of *Culex* spp. in the San Joaquin Valley (27). To briefly summarize MacDonald et al 2024, clusters of trap stations were created using a complete-linkage hierarchical clustering method with a tree height cutoff of ∼3,000 m using the ‘hclust’ function in R (R Core Team 2024). Each cluster contains distinct trap stations, so clusters of trap stations do not contain duplicate surveillance data, resulting in 96 unique clusters (Supplement Fig. 2). After clustering trap stations to 96 unique clusters (referred to as clusterID), we calculate monthly averages of 1) number of mosquitos per trap night, and 2) minimum WNV infection rate (MIR) for each cluster. The MIR was calculated with the following equation:

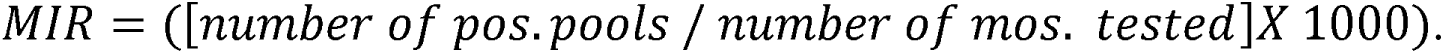

### Environmental and ecological covariates

Mosquito abundance and WNV risk are influenced by various factors, including the physiological development of individual mosquitos, the transmission dynamics of WNV, the quality of larval breeding sources, and the availability of habitats for adult mosquitos and their blood hosts (6,7,18) (Fig. 2). To isolate the impact of drought on mosquito abundance and WNV infection rates, we use the following set of covariates in our analysis: Palmer Drought Severity Index (PDSI), Kern River discharge rates, and bird community competence index. Data is summarized by clusterID within the 1,500 m radius buffer around trap station cluster centroids. Covariates in our models are standardized.

**Figure 2.**
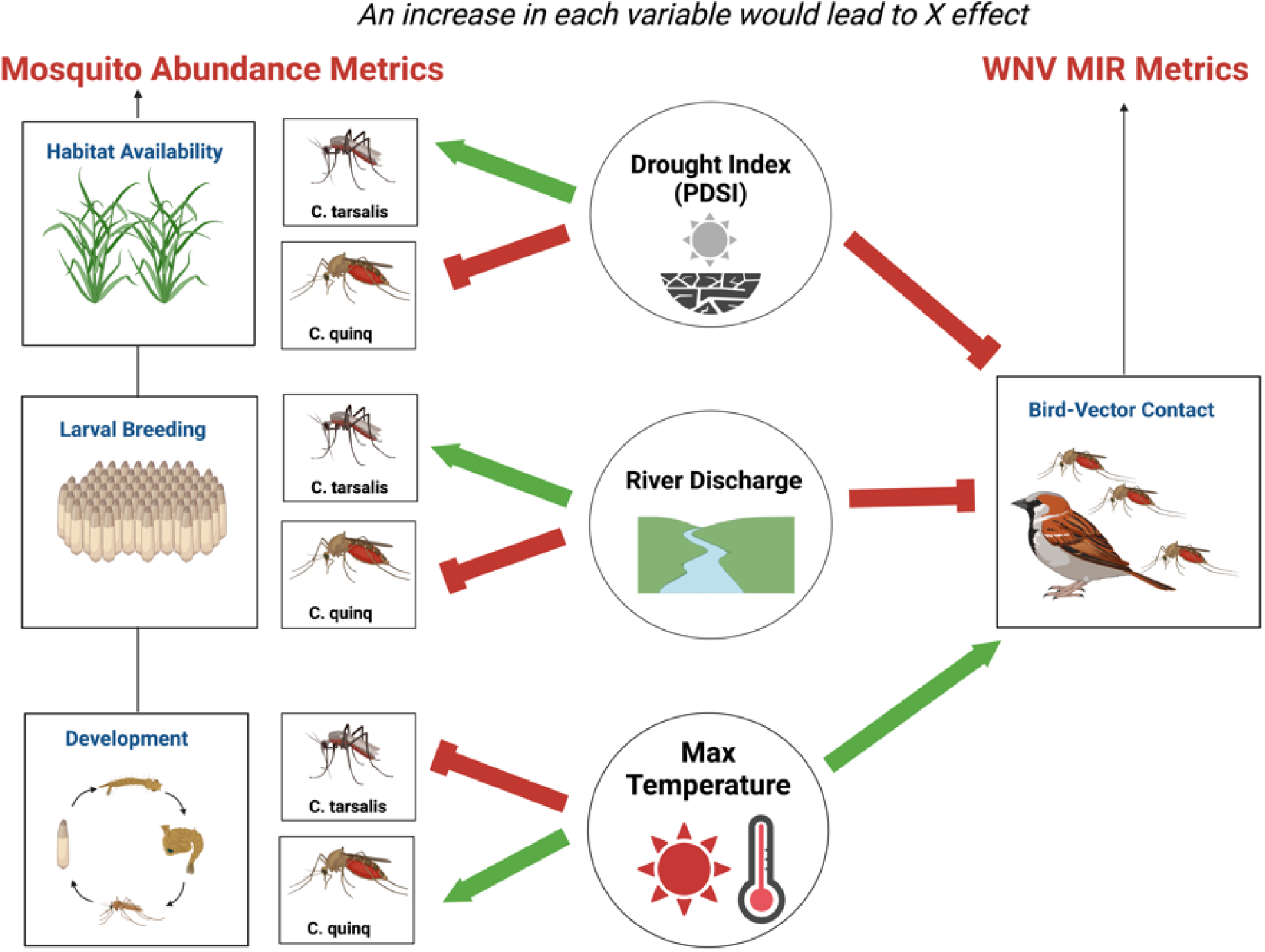
Hypothesized relationship between drought measurements, adult *Culex* spp. mosquito abundances and West Nile virus minimum infection rates (WNV MIR). Changes in a drought measurement (circle) can either negatively (red) or positively (green) affect mosquito-abundance or bird-mosquito contact rates, influencing WNV dynamics. The Palmer Drought Severity Index (PDSI), which ranges from 4 (wet) to −4 (dry), indicates wetter conditions with higher values. Image created with BioRender.

To quantify drought conditions throughout our study period, we use the standardized Palmer Drought Severity Index (PDSI) from gridMET, that estimates the relative soil moisture conditions every five days. A PDSI value > 4 represents very wet conditions, while a PDSI value < −4 represents extreme drought (29). We average the PDSI index per time step (month-year) and clusterID (Fig. 3). We hypothesize that a positive PDSI value represents a landscape more suitable for larval breeding, with sufficient vegetation shelter for adults and their blood meal hosts, compared to a negative PDSI value. We also include a month-lag PDSI value to represent drought conditions in the prior month assuming that a wetter prior month would be more conducive to mosquito breeding and influence adult mosquito abundance in the current month, in contrast to a drier prior month.

**Figure 3.**
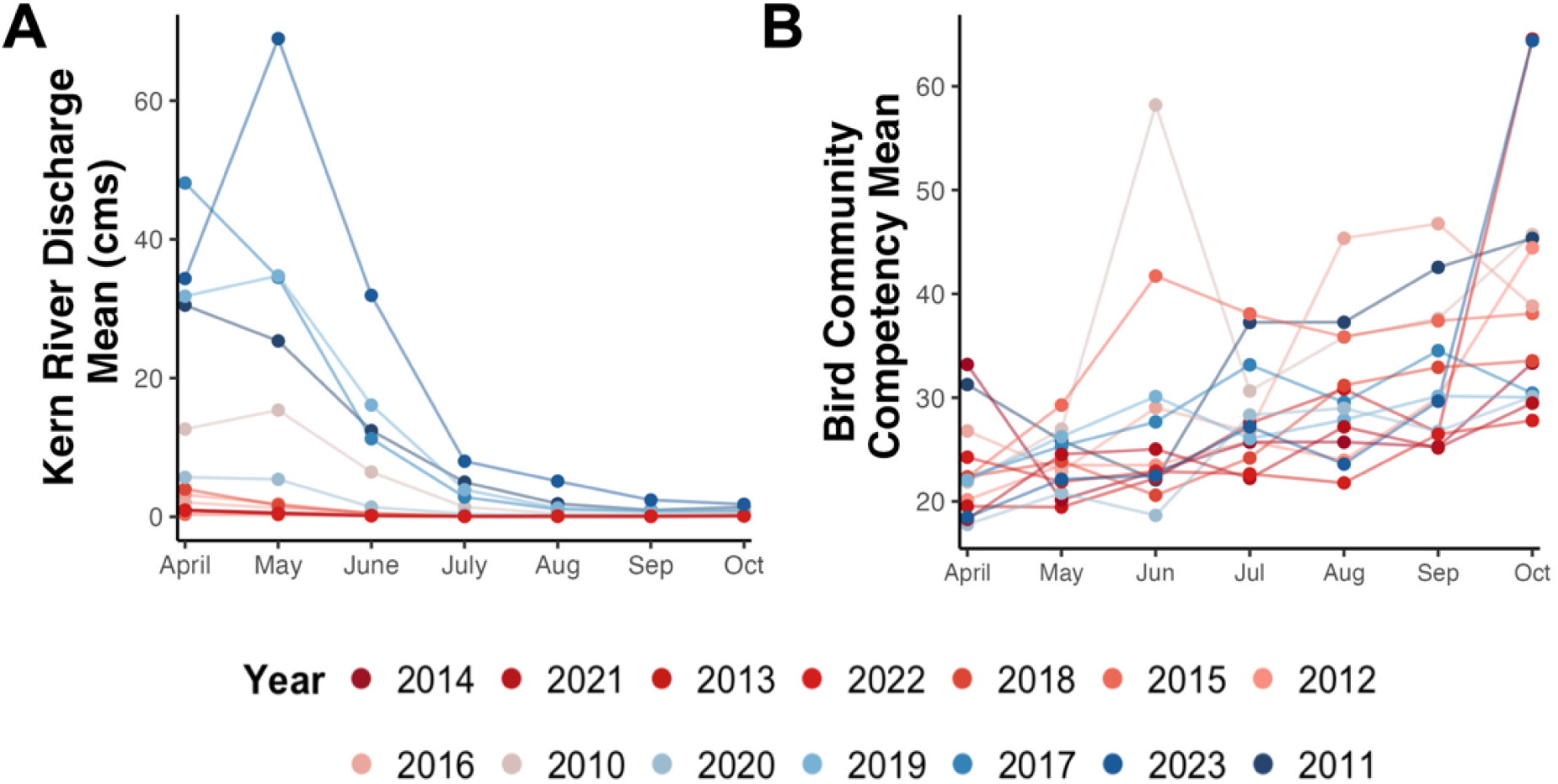
Monthly and yearly variation of environmental variables affecting (A) larval breeding water availability, and (B) likelihood of mosquitos taking an infectious bird blood meal. Colors represent the mean Palmer Drought Severity Index (PDSI) for each year, from the driest (−4.2 in 2014) to the wettest (4.5 in 2011).

To capture potential effects of water input into the system that are not captured by drought conditions (i.e., PDSI index) we included the monthly average discharge rate of the Kern River (cms) that is the same for all clusterIDs per month-year (16). The discharge rate data was sourced from the USGS NWIS for station SF Kern R NR Onyx CA - 11189500 (latitude: 35.7374516, longitude: −118.173689). The Kern River watershed is a dynamic source of water (Fig. 3) that is dependent on the Sierra Nevada snowpack, with water released from Lake Isabella tailored to meet local municipal and agricultural needs (12,14). During years of above-normal rainfall, the Kern River flowing through Bakersfield causes intentional flooding of ponds to recharge aquifers and directs excess flow into Lake Buena Vista, located east of Bakersfield (12,14). Previous studies have found a correlation between increased discharge rates of Kern River and *Cx. tarsalis* abundance (15). We hypothesize that during years with relatively high Kern River discharge rates, there will be more standing water in the surrounding areas of Bakersfield, which will increase available breeding habitat for *Cx. tarsalis*, increasing their abundances. Whereas *Cx. quinquefasciatus* breeds in smaller bodies of water (i.e., containers and storm drains) so an increase in Kern River discharge rate may flush out *Cx. quinquefasciatus* larvae that are breeding in the portions of the Kern River that flow through the city of Bakersfield.

In addition to climate covariates, birds are generally understood to be key ecological drivers of WNV transmission dynamics, serving as amplification hosts of vector-borne diseases. Thus, it is important to incorporate the abundance of birds in the study area and consider their species-specific relative competencies as potential hosts of WNV, as not all birds that migrate through or are residents of Kern County are competent hosts for West Nile virus (19). Building upon MacDonald et al 2024, we created a bird community competence index that we average by month-year for each clusterID (Fig. 3). Specifically, the relative abundances of 38 bird species inhabiting the Central Valley were modeled using *Best Practices for Using eBird Data* adopted from Cornell Ornithology Laboratory to replicate eBird status and trends products (30) (Supplemental Table 1). Further details can be found in the Supplement Text 1. We hypothesize that months with higher community competence resulted in a higher probability of a mosquito taking an infectious blood meal, resulting in a higher WNV minimum infection rate (MIR).

### Isolating the mosquito-borne disease risk and drought relationships

Randomized, controlled experiments are the gold standard for inferring causal effects in ecological communities (31,32). However, in observational studies involving complex ecological systems, it is often infeasible to randomly assign treatment due to resource limitations or ethical concerns (31). Here we use a within-estimator panel model (also known as a “fixed effects” model in econometrics) to approximate randomized experiments in observational data settings (31,32).

Mosquito-borne disease risk is impacted by multiple abiotic and biotic factors; however, our focus is the impact of drought on mosquito abundance and WNV infection rates. To parse drought from other relevant factors, we use a within-estimator model with time (i.e., month) and individual (i.e., clusterID) fixed effects in addition to key time-varying biotic and abiotic covariates hypothesized to influence our outcome variables of interest. Key time-varying variables such as Kern River discharge rate and bird community competence influence mosquito abundance and mosquito-bird contact rates, which can impact WNV transmission rates. Our goal is to account for or remove, through differencing, all variables that are correlated with our covariate of interest (i.e., drought severity; PDSI) and our outcomes, as any such variable that remains in the error term would bias our estimation. Variables that drive the outcome, but are not correlated with drought would increase noise, but would not bias our coefficient estimates.

Our final mosquito abundance model predicts the number of mosquitos per trap night in clusterID *i* and month *t* (log(*Abundance_it_*)) with the drought severity from the previous month (*PDSI_it_*_-1_) and is written as

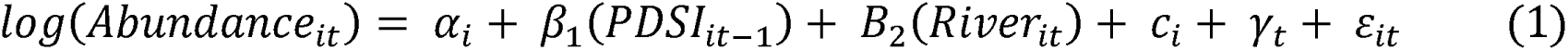

We control for average Kern River discharge rates of the contemporaneous month and include the fixed-effect dummy variables clusterID (*c_i_*) and month (γ*_t_*). We include the terms *α_i_* for clusterID intercept and *ε_it_* as the random error. We run separate models for each focal species, *Cx. tarsalis* and *Cx. quinquefasciatus*.

Our final mosquito infection model predicts West Nile virus minimum infection rate in clusterID *i* and month *t* (log(*MIR_it_*)) with drought severity index from the previous month (*PDSI_it_*_-1_) and is written as

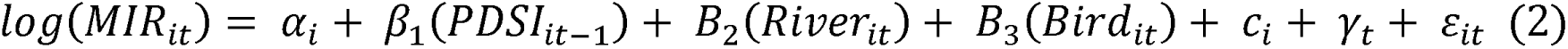

We additionally control for average Kern River discharge rates of the contemporaneous month and the average bird host community competence of the contemporaneous month and include the fixed-effect dummy variables clusterID (*c_i_*) and month (γ*_t_*). We include the terms *α_i_* for clusterID intercept and *ε_it_* as the random error. We run separate models for each focal species, *Cx. tarsalis* and *Cx. quinquefasciatus*, infection rates.

We estimated the within-estimator panel models using the feols function from the ‘fixest’ package (Bergé 2018). For our final models, we calculate Conley standard errors to account for spatial autocorrelation of the standard errors (see Supplement Text 2 for further model justifications).

All statistical analyses were performed using R software version 4.4.0 (RStudio Core Team 2024). Data cleaning and visualizations were conducted with ‘tidyversè and ‘ggplot2’ packages, respectively (34,35). Code for figure making and the panel model analysis can be found here: https://github.com/sbsambado/wnv_drought_kern.

## RESULTS

A total of 2,353,573 *Cx. tarsalis* (per trap night mean = 14.0 ± 82.1) and 1,321,196 *Cx. quinquefasciatus* (per trap night mean = 9.3 ± 35.0) adult female mosquitos were captured from April through October between 2010-2023 in Kern County (Table 1). Of those mosquitos captured and tested, *Cx. tarsalis* and *Cx. quinquefasciatus* have a mean minimum WNV infection rate of 3.2 (± 9.5) and 4.1 (± 10.5), respectively (Fig. 2).

### Culex spp. mosquito abundances are impacted by drought severity

We found a significant effect of drought severity on the abundances of both species of mosquito. However, the magnitude of the effect differed between mosquito species (Fig. 4). A higher value of the drought index means wetter conditions; thus, our results can be interpreted as one standard deviation increase in wet conditions leads to a 15% increase in *Cx. tarsalis* abundance (beta = 0.15 ± 0.041). We did not find a significant effect for *Cx. quinquefasciatus* abundance (beta = 0.018 ± 0.030), but it was positively trending similarly to *Cx. tarsalis* (Table 2). As expected, the control covariate for Kern River discharge rate was positive for both mosquito species but had a larger impact on *Cx. tarsalis* compared to *Cx. quinquefasciatus*. This suggests that our panel models are effectively isolating the effect of drought on mosquito abundances, independent of the water pulses into the system not captured by drought metrics.

**Figure 4.**
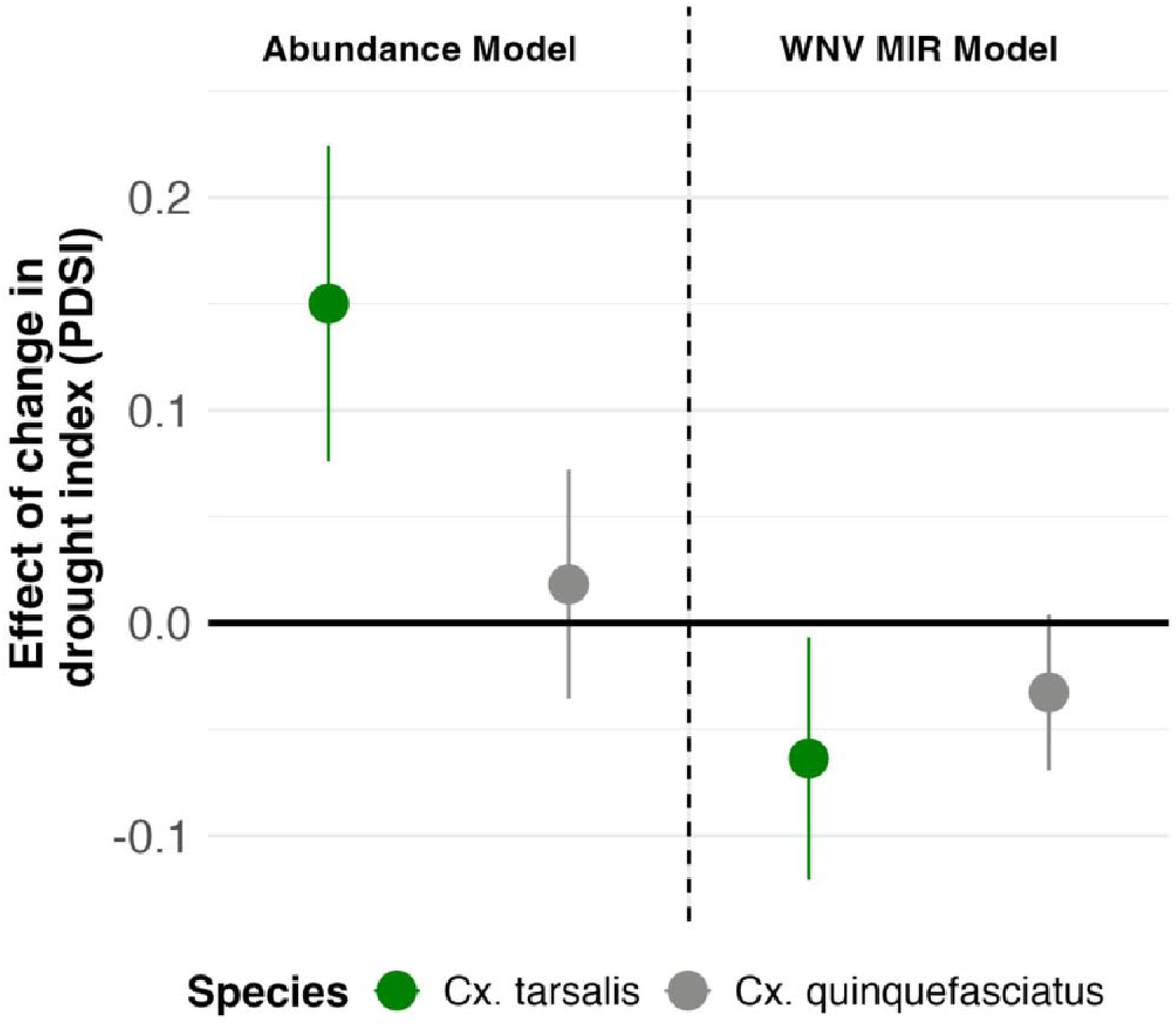
Coefficient estimates and 95% confidence intervals for panel models of the log_10_ transformed mosquito abundances and WNV minimum infection rates (WNV MIR), by species. An increase in PDSI represents an increase in wetness.

**Table 2.** Results for the preferred model specification best fit for the abundance and WNV minimum infection rate (MIR) models for both species of mosquitos, *Cx. tarsalis* and *Cx. quinquefasciatus*, with the fixed effects of clusterID and month *t*. Conley standard errors were used to account for spatial correlation of the standard errors. All covariates are the monthly average standardized.

|  |  | *Estimate* | *Conley SE* | *t-value* | *p-value* |
| --- | --- | --- | --- | --- | --- |
| **ABUNDANCE** | **C. tarsalis** |  |  |  |  |
| **MODELS** | PDSI<sub>t-1</sub> | 0.15 | 0.04 | 3.67 | < **0.001** |
|  | River Discharge<sub>t</sub> | 0.59 | 0.05 | 11.76 | < 0.001 |
|  | **C. quinquefasciatus** |  |  |  |  |
|  | PDSI<sub>t-1</sub> | 0.018 | 0.03 | 0.60 | **0.55** |
|  | River Discharge<sub>t</sub> | 0.11 | 0.02 | 4.56 | < 0.001 |
| **WNV MIR** | **C. tarsalis** |  |  |  |  |
| **MODELS** | PDSI<sub>t-1</sub> | -0.064 | 0.01 | -4.80 | < **0.001** |
|  | River Discharge<sub>t</sub> | -0.029 | 0.005 | -5.85 | < 0.001 |
|  | Bird community<sub>t</sub> | 0.090 | 0.04 | 2.21 | 0.027 |
|  | **C. quinquefasciatus** |  |  |  |  |
|  | PDSI<sub>t-1</sub> | -0.032 | 0.01 | -2.29 | **0.022** |
|  | River Discharge<sub>t</sub> | -0.036 | 0.007 | -5.49 | < 0.001 |
|  | Bird community<sub>t</sub> | 0.042 | 0.03 | 1.50 | 0.013 |

### WNV infection rates are impacted by drought severity

We found a significant effect of drought on the West Nile infection rate of both mosquito species (Fig. 4). A one standard deviation increase in wet conditions of the prior month leads to a 6% and 3% decrease in WNV MIR infection of *Cx. tarsalis* (beta = −0.064 ± 0.013) and *Cx. quinquefasciatus* (beta = −0.033 ± 0.014), respectively. (Table 2). The control covariates for Kern River discharge rate and bird community competence were trending in the expected directions (negative for Kern River, positive for bird competence) further suggesting our within-estimator panel models are isolating the effect of drought on WNV minimum infection rates, independent of pulses of water into the system not captured by drought metrics and controlling for changes in bird host community competence.

## DISCUSSION

Climate change is driving an increase in both the frequency and severity of droughts (10,11). In Mediterranean climates like California, this is resulting in abrupt shifts between periods of intense droughts and severe atmospheric river and precipitation events, profoundly impacting entire ecological communities (10,24). Drought exerts both direct and indirect influences on the risk of mosquito-borne diseases (5,7). We find that drought is reducing mosquito abundances while concurrently increasing West Nile virus infection rates. However, the extent to which drought affects the abundance of two key WNV vectors, *Cx. tarsalis* and *Cx. quinquefasciatus*, varies. We hypothesize that this variation is due to their distinct ecological niches, which directly influence their breeding preferences and responses to drought conditions. Drought may indirectly influence WNV infection rates by reducing bodies of water on the landscape, potentially intensifying mosquito-bird contact rates and thereby contributing to increased WNV infection rates (7,14). Here, we present evidence of contrasting effects of drought on two critical metrics of mosquito-borne disease, but the effects differ from one vector species to another suggesting within the same period, human health risks will vary from rural to urban settings. Our findings underscore the importance of species-specific responses to drought, carrying significant implications for public health messaging and resource allocations for control measures.

The risk of mosquito-borne diseases depends on both mosquito abundance and infection rates – two outcomes that are affected by drought but respond in distinct ways. Our findings revealed that after a dry period in the previous month (indicated by a lower PDSI index), mosquito abundances decreased while West Nile virus infection rates increased. These findings align with our hypotheses and support previous studies (7,12,14,25), which suggest that drought reduces available mosquito breeding habitats, thereby lowering population numbers. Drought not only reduces suitable breeding habitat but overall area of standing water on a landscape, potentially leading to greater aggregation of mosquitos and competent bird hosts, increasing contact rates and thus WNV transmission (7,14). However, the average magnitude of the effect of drought on WNV is smaller compared to our biological control – bird community competence index – suggesting that other factors are also influencing WNV infection rates (18,36,37). As expected, we find a positive relationship between WNV infection rates and bird community competence. As California experiences more dramatic shifts in drought severity, mosquito-borne disease risk may similarly be increasingly amplified and dampened in response. However, future research should investigate how prolonged severe droughts affect transmission potential and determine if there are thresholds in the response of mosquitos and bird hosts that might decouple mosquito-borne disease risk from drought conditions.

The impact of drought on mosquito abundance differs between the two key vector species in Kern County, California. Our findings show that drought has a more pronounced effect on the rural vector, *Cx. tarsalis*, compared to the urban vector, *Cx. quinquefasciatus*. This suggests that understanding species-specific habitat and behavioral traits is crucial for understanding the impact of drought on mosquito ecology (8,12,25). The rural mosquito, *Cx. tarsalis*, which is commonly found in agricultural areas, appears to be more sensitive to drought fluctuations, possibly because it has less refugia than the urban mosquito during unfavorable drought conditions (12,14). While *Cx. quinquefasciatus*, the urban mosquito, was negatively affected by drought, its abundances exhibited less variation in response to fluctuating drought conditions than the rural mosquito (Reisen 2009). This resilience may stem from the mosquito’s ability to seek refuge and breed in human-made containers such as swimming pools, ornamental plants, and urban trash (36). As expected in our mosquito abundance models, our control for the Kern River discharge rate revealed a greater impact on the rural vector than the urban vector. This difference may be attributed to breeding preferences: an increase in Kern River discharge positively affects rural mosquitos that thrive with large water sources (e.g., flooded aquifers, seasonal wetlands), whereas urban mosquitos, which typically breed in smaller containers, are negatively impacted by an increase in discharge rates because larvae are flushed from their habitats (12,15,16).

While our study identifies contrasting effects of drought on West Nile virus risk that vary across vector species, it is essential to consider these results in the context of the Central Valley. In both rural and urban settings, humans modify water availability through irrigation and landscaping practices. Even during drought years, substantial federal and local funding supports the extensive agricultural irrigation networks that can supplement natural breeding habitats. Regions in California, such as Sacramento, Bakersfield, and Coachella Valley, are endemic for WNV but differ in their water management practices, which can complicate predicting WNV risk in response to drought (8,14). Sacramento and Bakersfield benefit from natural waterways (i.e. Sacramento and Kern rivers), especially the Sacramento-San Joaquin River Delta, which is less prone to drying out during drought years (8). In contrast, Coachella Valley relies more heavily on human-driven irrigation rather than natural waterways, potentially altering the impact of drought on mosquito abundances and WNV risk depending on human behavior. Without the natural supplement of water, the effect of drought on West Nile risk may be greater by increasing infection rates in mosquitos, or if natural water bodies are robust to drought, then mosquito populations will remain stable. Another factor that could influence our mosquito abundance results is the application of pesticides by vector control agencies and private individuals. Some preliminary data suggests that due to the consistency of pesticide application by Kern Mosquito and Vector Control District staff, there would be little effect on our general results that encompass 14 years of surveillance data (Supplement Fig. 3). Another aspect that warrants further investigation is the impact of land cover type on our results. With the within-estimator panel models we aimed to remove variation that is unique to individual trap stations (31,32), but there could be further investigation due to future land use change as a result of projected human population growth in California’s Central Valley or the Sustainable Groundwater Management Act, which may lead to the fallowing of large areas currently in agricultural production (Public Policy Institute of California).

Evaluating mosquito-borne disease risk in response to drought must also account for areas where human populations are likely to encounter mosquitos (20–22). In our study system, we observe varying magnitudes of response to drought between rural and urban mosquitos, highlighting important considerations for public health messaging. During wet years, the rural mosquito *Cx. tarsalis* tends to experience a greater increase in abundance compared to the urban mosquito *Cx. quinquefasciatus*. This finding suggests that agricultural workers, who may have limited access to public health information or inequitable access to health care, are potentially at a higher risk of West Nile virus compared to individuals primarily spending time in urban areas (14,22,23). In dry years, when rural areas pose lower risk, mosquito mitigation efforts should concentrate on urban settings, particularly in areas adjacent to Kern River or residential pools, which could serve as refuges for mosquito populations (36,37). The differential responses of rural and urban mosquitos to drought underscores the need for targeted public health strategies that account for varying risk levels across different environments and human populations.

## CONCLUSION

Our study investigated the effect of drought severity on a complex vector-borne disease with a multi-vector system. We found that 1-month lagged drought conditions can exert contrasting effects on mosquito abundance, infection rates and ultimately mosquito-borne disease risk. Increased drought severity negatively affects mosquito abundances directly. In contrast, West Nile infection rates are positively influenced by increased drought severity, likely due to reduced water availability concentrating mosquito vectors and bird hosts, increasing vector-host contact rates and thereby increasing transmission risk. Importantly, the effects of drought vary between vector species, *Cx. tarsalis* (rural) and *Cx. quinquefasciatus* (urban), underscoring the need for nuanced public health messaging and mitigation strategies as drought conditions fluctuate in California.

## DATA AVAILABILITY STATEMENT

Upon acceptance, data, data analyses, code, and figures will be provided via GitHub (https://github.com/sbsambado/wnv_drought_kern) and stored on DataDyrad repository. The code is not novel. Details on data sources used for the analyses can be found in Supplemental Table 2. Additional data sources used throughout the main text can be found at: US Census Bureau https://data.census.gov/profile/Bakersfield_city,_California?g=160XX00US0603526; US Geological Survey National Water Information System (USGS NWIS) https://waterdata.usgs.gov/monitoring-location/11189500/#parameterCode=00065&period=P7D&showMedian=false; US Department of Agriculture National Agricultural Statistics Services (USDA NASS) https://www.nass.usda.gov/Publications/AgCensus/2022/Online_Resources/County_Profiles/California/cp06029.pdf; Public Policy Institute of California https://www.ppic.org/wp-content/uploads/content/pubs/jtf/JTF_CentralValleyJTF.pdf.

## ACKNOWLEDGMENTS

We acknowledge Kern Mosquito and Vector Control District, and specifically La Thao and Mark Dery, for providing surveillance data. We thank Nickolas McManus for providing feedback on the ideas behind this paper. SS wishes to acknowledge the foundational empirical research conducted by Dr. William Reisen, which significantly inspired and informed much of this work.

## FUNDING INFORMATION

A.J.M., A.L., D.S., & A.Q. acknowledge funding from the USDA National Institute of Food and Agriculture Rapid Response to Extreme Weather Events Across Food and Agricultural Systems program (Grant #2023-68016-40683). SS and AJM acknowledges funding support from the Training Grant Program of the Pacific Southwest Regional Center of Excellence for Vector-Borne Diseases funded by the U.S. CDC (Cooperative Agreement 1U01CK000516 and U01CK000649). A.L., D.S., & A.Q. acknowledge the USDA NIFA Sustainable Agroecosystems program (Grant #2022-67019-36397). A.J.M. further acknowledges the National Science Foundation and Fogarty International Center (DEB-2011147; DEB-2339209), and a Faculty Research Grant from the UCSB Academic Senate. D.S. gratefully acknowledges the NASA EMIT Science and Applications Team Program (Grant #80NSSC24K0861), the NASA Land-Cover/Land Use Change program (Grant #NNH21ZDA001N-LCLUC), the NASA Remote Sensing of Water Quality program (Grant #80NSSC22K0907), the NASA Applications-Oriented Augmentations for Research and Analysis program (Grant #80NSSC23K1460), the NASA Commercial Smallsat Data Analysis program (Grant #80NSSC24K0052), the NASA FireSense Airborne Science Program (Grant #80NSSC24K0145), the California Climate Action Seed Award Program, and the NSF Signals in the Soil program (Award #2226649).

## Statement of authorship

**SS:** Conceptualization; data curation; formal analysis; visualization; writing – original draft; writing – review and editing. **TS:** data curation; writing – review and editing. **ZR:** data curation; methodology; writing – review and editing. **AL:** funding acquisition; methodology; supervision; writing – review and editing. **JC:** data curation; visualization; writing review and editing. **AQ:** funding acquisition; writing – review and editing. **DS:** funding acquisition; writing – review and editing. **AJM:** conceptualization; funding acquisition; project administration; resources; supervision; writing – review and editing.

## Notes

### Competing Interest Statement

The authors have declared no competing interest.

