## Supplemental Information for "The paradoxical impact of drought on West Nile virus risk: insights from long-term ecological data"

**SUPPORTING INFORMATION**

**Supplemental Text**

S Text 1. Detailed methods on bird community competence calculation

S Text 2. Detailed methods on within-estimator panel models

**Supplemental Tables**

S Table 1. List of bird species used for bird community competence calculations

S Table 2. Summary of data sources

S Table 3. Model summaries for *Cx. tarsalis* abundance

S Table 4. Model summaries for *Cx. quinquefasciatus* abundance

S Table 5. Model summaries for *Cx. tarsalis* WNV MIR

S Table 6. Model summaries for *Cx. quinquefasciatus* WNV MIR

**Supplemental Figures**

S Figure 1. Drought conditions in Kern County between 2010-2023

S Figure 2. Study site trap station and clusterID information

S Figure 3. Pesticide application in Kern County

S Figure 4. Coefficient estimates for abundance models with different fixed effects

S Figure 5. Coefficient estimates for WNV MIR models with different fixed effects

S Figure 6. Coefficient estimates for abundance models with different Conley standard errors

S Figure 7. Coefficient estimates for WNV MIR models with different Conley standard errors

S Figure 8. Coefficient estimates for abundance models with different types of standard errors

S Figure 9. Coefficient estimates for WNV MIR models with different types of standard errors

**Supplemental References**

**Supplemental Text 1. Detailed methods on bird community competence calculation**

In addition to climate covariates, birds are generally understood to be key ecological drivers of WNV transmission dynamics, serving as amplification hosts of vector-borne diseases. Thus, it is important to incorporate the abundance of birds in the study area and consider their species-specific relative competencies as potential hosts of WNV, as not all birds that migrate through or are residents of Kern County are competent hosts (Kilpatrick et al. 2007). Building upon MacDonald et al. 2024, we created a bird community competence index that we average by month-year for each clusterID. Specifically, the relative abundances of 38 bird species inhabiting California’s Central Valley were modeled using *Best Practices for Using eBird Data* adopted from Cornell Ornithology Laboratory to replicate eBird status and trends products (Fink et al. 2022). Raw bird observation data was sourced from the eBird Basic Dataset and was processed in R (Strimas-Mackey et al. 2023). The NASA MODIS land cover product “MCD12Q1” (Friedl et al. 2022) and elevation pulled from EarthEnv.org (Amatulli et al. 2018) were used to create a prediction grid for the species distribution model. Encounter rate was first modeled to give a proportional estimate of occupancy probability, or the probability that a species occurs in a given area. These modeled counts were then used to create species-specific weekly relative abundance models, expressed as an index of the expected count of individuals of a species in a given area. Random forest machine learning was used to estimate both encounter rate and relative abundance and was modeled to species-specific raster stacks. Due to data sparsity for modeling bird relative abundance for every week, the modeling process was conducted across the extent of the Central Valley. As not every species was represented in the eBird data for every week of the study period, and could thus not be modeled for every week, linear interpolation was used to fill out missing raster layers. This was done by generating empty raster layers for unmodeled periods and using the `threadr` package to interpolate empty layers in between modeled layers and extrapolate trailing and leading empty layers (Grange et al. 2024). To handle species which were too sparsely modeled to even be interpolated, zero-filled rasters were created to fill missing time periods. These complete weekly species rasters were then stacked by year and multiplied by the corresponding family-average community competence index to obtain yearly competence rasters for every species (Kilpatrick et al. 2007). These species-specific yearly competence rasters were summed together between species using the `terra` package in R to obtain yearly community competency rasters layered by week (Hijmans et al. 2024). To calculate zonal statistics for the mean weekly community competency of each trap station cluster,`terra` was used using polygons of the Kern County trap station clusters as zones, averaged by month-year.

**Supplemental Text 2.** **Detailed methods on within-estimator panel models**

Randomized, controlled experiments are the gold standard for inferring causal effects in ecological communities (Larsen et al. 2019, Dee et al. 2023). However, in observational studies involving complex ecological systems, it is often infeasible to randomly assign treatment due to resource limitation or ethical concerns (Larsen et al. 2019). Here we use a within-estimator panel model (also known as a “fixed effects” model in econometrics) to approximate randomized experiments in observational data settings (Larsen et al. 2019, Dee et al. 2023). To infer plausibly causal effects of drought severity (PDSI) on mosquito-borne disease risk – mosquito abundances and WNV infection rates – we addressed some confounding variables that may introduce bias to our results. Some confounding variables may not vary over the study period (i.e., time-invariant) and are unique to each individual clusterID such as distance to Kern River. Other confounding variables that do vary over the study period but are shared by all clusterIDs (i.e., time-shocks) would be budget cuts to surveillance efforts which could have unobserved effects on our results. By controlling for individual and time fixed effects, as well as leveraging longitudinal data from repeated samples of the same clusterIDs over time, we remove variation in our outcome attributed to confounding variables, hence better isolating the effect of drought on mosquito-borne disease risk. With this approach, we can look at clusterID-specific means and estimate the deviation from the clusterID-specific mean as a function of time-varying explanatory variables.

Supplemental Tables and Figures below highlight the model building process of varying fixed effect terms, varying distances for Conley standard errors, and varying standard error calculations.

**Supplemental Tables**

**Table S1. List of bird species used for bird community competence calculations**

Table of 38 bird species inhabiting the Central Valley that were modeled using *Best Practices for Using eBird Data* adopted from Cornell Ornithology Laboratory to replicate eBird status and trends (Fink et al. 2022).

| **Scientific name** | **Common name** |
| --- | --- |
| *Anas platyrhynchos* | mallard |
| *Agelaius phoeniceus* | red-winged blackbird |
| *Agelaius tricolor* | tricolored blackbird |
| *Aphelocoma californica* | California scrub-jay |
| *Aphelocoma woodhouseii* | Woodhouse's scrub-jay |
| *Branta canadensis* | canada goose |
| *Bubo virginianus* | great horned owl |
| *Bubulcus ibis* | cattle egret |
| *Buteo jamaicensis* | red-tailed hawk |
| *Callipepla californica* | Galifornia quail |
| *Callipepla gambelii* | Gambel's quail |
| *Catharus ustulatus* | Swainson's thrush |
| *Charadrius vociferus* | killdeer |
| *Colaptes auratus* | northern flicker |
| *Colinus virginianus* | northern bobwhite |
| *Columba livia* | rock pigeon |
| *Columbina passerina* | common ground dove |
| *Corvus brachyrhynchos* | American crow |
| *Cyanocitta cristata* | blue jay |
| *Dumetella carolinensis* | gray catbird |
| *Euphagus cyanocephalus* | Brewer's blackbird |
| *Falco sparverius* | American kestrel |
| *Fulica americana* | American coot |
| *Haemorhous mexicanus* | house finch |
| *Hylocichla mustelina* | wood thrush |
| *Larus delawarensis* | ring-billed gull |
| *Melopsittacus undulatus* | budgerigar |
| *Melospiza melodia* | song sparrow |
| *Mimus polyglottos* | northern mockingbird |
| *Molothrus ater* | brown-headed cowbird |
| *Nycticorax nycticorax* | black-crowned night heron |
| *Passer domesticus* | house sparrow |
| *Phasianus colchicus* | ring-necked pheasant |
| *Quiscalus quiscula* | common grackle |
| *Sturnus vulgaris* | European starling |
| *Turdus migratorius* | American robin |
| *Tyto alba* | barn owl |
| *Zenaida macroura* | mourning dove |

**Table S2. Summary of data sources**

Summary of data sources used in the main and supplemental files with respective hyperlinks and references (below). All data (except Kern River discharge rates) were summarized to each clusterID and averaged per month. Mosquito surveillance data was only adult, female *Culex tarsalis* and *Culex quinquefasciatus* captured between April through October between 2010-2023.

| **Type** | **Data** | **Source** | **Hyperlink** |
| --- | --- | --- | --- |
| **Vector** | Mosquito abundance | KMVCD | NA |
|  | Mosquito WNV infection | KMVCD | NA |
| **Host** | Bird community competence | eBird^1^ | <https://science.ebird.org/en/status-and-trends/download-data> |
| **Environmental** | Temperature (minimum/maximum) | gridMET^2^  (~ 4km; daily) | <https://www.drought.gov/data-maps-tools/gridded-surface-meteorological-gridmet-dataset> |
|  | Precipitation (mm) | gridMET^2^  (~ 4km; daily) | <https://www.drought.gov/data-maps-tools/gridded-surface-meteorological-gridmet-dataset> |
|  | PDSI (index -4 to 4) | gridMET^2^  (~ 4km; weekly) | <https://www.drought.gov/data-maps-tools/us-gridded-palmer-drought-severity-index-pdsi-gridmet> |
|  | Land Cover Type (MCD12Q1) | MODIS^3^ | <https://lpdaac.usgs.gov/products/mcd12q1v061/> |
|  | Elevation | EarthEnv | <https://www.earthenv.org/topography> |
|  | River discharge (cms) | USGS NWIS  (SF Kern R NR Onyx CA - 11189500 station, weekly) | <https://waterdata.usgs.gov/monitoring-location/11189500/#parameterCode=00065&period=P7D&showMedian=false> |
|  | 30-year normal | PRISM | <https://prism.oregonstate.edu/normals/> |
|  | Land Cove Type  (S Fig. 2B) | FFMP | <https://www.conservation.ca.gov/dlrp/fmmp> |

References: ^1^Fink et al. 2022, ^2^Abatzoglou 2013, ^3^Friedl 2022

**Table S3. Model summaries for *Cx. tarsalis* abundance.** The effects of environmental and ecological covariates on the log_10_ transformed average **monthly abundance** of adult female *Cx. tarsalis*. Each column represents a different model (1-6), where the included fixed effects are denoted by ‘X’. Estimate (error). All covariates are standardized. Unless otherwise noted, we use Conley standard errors (11 km). For all models, p < 0.001***, p < 0.01**, p < 0.05*.

| **VARIABLES** | **(1)** | **(2)** | **(3)** | **(4)** | **(5)** | **(6)** |
| --- | --- | --- | --- | --- | --- | --- |
| **PDSI**  *(prior month)* | 0.29  (0.05)  *** | 0.30  (0.06)  *** | 0.14  (0.04)  *** | 0.15  (0.2) | 0.15 (0.04) *** | -0.12  (0.2) |
| **River Discharge**  (*current month)* | 0.33  (0.1)  ** | 0.36  (0.1)  ** | 0.57  (0.06)  *** | 0.19  (0.1) | 0.59 (0.05) *** | 0.45 (0.04)  *** |
| **ClustID** |  | X |  |  | X | X |
| **Month** |  |  | X |  | X | X |
| **Year** |  |  |  | X |  | X |
| **Observations** | 5,076 | 5,076 | 5,076 | 5,076 | 5,076 | 5,076 |

**Table S4. Model summaries for *Cx. quinquefasciatus* abundance.** The effects of environmental and ecological covariates on the log_10_ transformed average **monthly abundance** of adult female *Cx. quinquefasciatus*. Each column represents a different model (1-6), where the included fixed effects are denoted by ‘X’. Estimate (error). All covariates are standardized. Unless otherwise noted, we use Conley standard errors (11 km). For all models, p < 0.001***, p < 0.01**, p < 0.05*.

| **VARIABLES** | **(1)** | **(2)** | **(3)** | **(4)** | **(5)** | **(6)** |
| --- | --- | --- | --- | --- | --- | --- |
| **PDSI**  *(prior month)* | 0.17  (0.04)  *** | 0.17  (0.03)  *** | 0.016  (0.04) | 0.53  (0.1)  *** | 0.018 (0.03) | -0.10  (0.05) |
| **River Discharge**  (*current month)* | -0.15  (0.03)  *** | 0.15  (0.03)  *** | 0.11  (0.02)  *** | -0.2  (0.04)  *** | 0.11 (0.02) *** | 0.09  (0.02)  *** |
| **ClustID** |  | X |  |  | X | X |
| **Month** |  |  | X |  | X | X |
| **Year** |  |  |  | X |  | X |
| **Observations** | 5,445 | 5,445 | 5,445 | 5,445 | 5,445 | 5,445 |

**Table S5. Model summaries for *Cx. tarsalis* WNV MIR.** The effects of environmental and ecological covariates on the log_10_ transformed average **monthly WNV minimum infection rate (MIR)** of adult female *Cx. tarsalis*. Each column represents a different model (1-6), where the included fixed effects are denoted by ‘X’. Estimate (error). Unless otherwise noted, we use Conley standard errors (11 km). All covariates are standardized. For all models, p < 0.001***, p < 0.01**, p < 0.05*.

| **VARIABLES** | **(1)** | **(2)** | **(3)** | **(4)** | **(5)** | **(6)** |
| --- | --- | --- | --- | --- | --- | --- |
| **PDSI**  *(prior month)* | -0.14  (0.02)  *** | -0.13 (0.01)  *** | -0.08 (0.02)  *** | -0.18 (0.04) *** | -0.063 (0.01) *** | -0.023 (0.04) |
| **River Discharge**  (*current month)* | -0.17  (0.01)  *** | -0.16 (0.01)  *** | -0.02 (0.007)  ** | -0.22 (0.03) *** | -0.29 (0.005) *** | -0.059 (0.03) |
| **Bird Community Competence**  (*current month*) | 0.15  (0.04)  *** | 0.15  (0.05)  ** | 0.093  (0.4)  * | 0.14 (0.05)  ** | 0.090 (0.04)  * | 0.056 (0.03) |
| **ClustID** |  | X |  |  | X | X |
| **Month** |  |  | X |  | X | X |
| **Year** |  |  |  | X |  | X |
| **Observations** | 3,037 | 3,037 | 3,037 | 3,037 | 3,037 | 3,037 |

**Table S3. Model summaries for *Cx. quinquefasciatus* WNV MIR.** The effects of environmental and ecological covariates on the log_10_ transformed average **monthly WNV minimum infection rate (MIR)** of adult female *Cx. quinquefasciatus*. Each column represents a different model (1-14), where the included fixed effects are denoted by ‘X’. Estimate (error). All covariates are standardized. Unless otherwise noted, we use Conley standard errors (11 km). For all models, p < 0.001***, p < 0.01**, p < 0.05*.

| **VARIABLES** | **(1)** | **(2)** | **(3)** | **(4)** | **(5)** | **(6)** |
| --- | --- | --- | --- | --- | --- | --- |
| **PDSI**  *(prior month)* | 0.0047 (0.02) | 0.00014 (0.02) | -0.014 (0.01) | -0.063  (0.03)  * | -0.033  (0.01)  * | -0.055  (0.02)  * |
| **River Discharge**  (*current month)* | -0.15 (0.007)  *** | -0.15 (0.002)  *** | -0.003 (0.006)  *** | -0.27  (0.01)  *** | -0.036  (0.007)  *** | -0.10  (0.01)  *** |
| **Bird Community Competence**  (*current month*) | 0.12  (0.03)  *** | 0.13  (0.04)  *** | 0.068 (0.03)  * | 0.063  (0.03)  * | 0.042  (0.03) | 0.016  (0.02) |
| **ClustID** |  | X |  |  | X | X |
| **Month** |  |  | X |  | X | X |
| **Year** |  |  |  | X |  | X |
| **Observations** | 6,320 | 6,320 | 6,320 | 6,320 | 6,320 | 5,445 |

**Supplemental Figures**

**Figure S1. Drought conditions in Kern County between 2010-2023**. Kern County drought history between 2010-2023 for each clusterID: (A) Palmer Drought Severity Index (-4 dry to 4 wet), (B) maximum mean temperature, (B) total yearly precipitation per rain year (October-March). Gray lines in (B) and (C) represent the 30-year normals for Kern County from USGS Parameter-elevation Regressions on Independent Slopes Model data portal (PRISM, <https://prism.oregonstate.edu/normals/>).

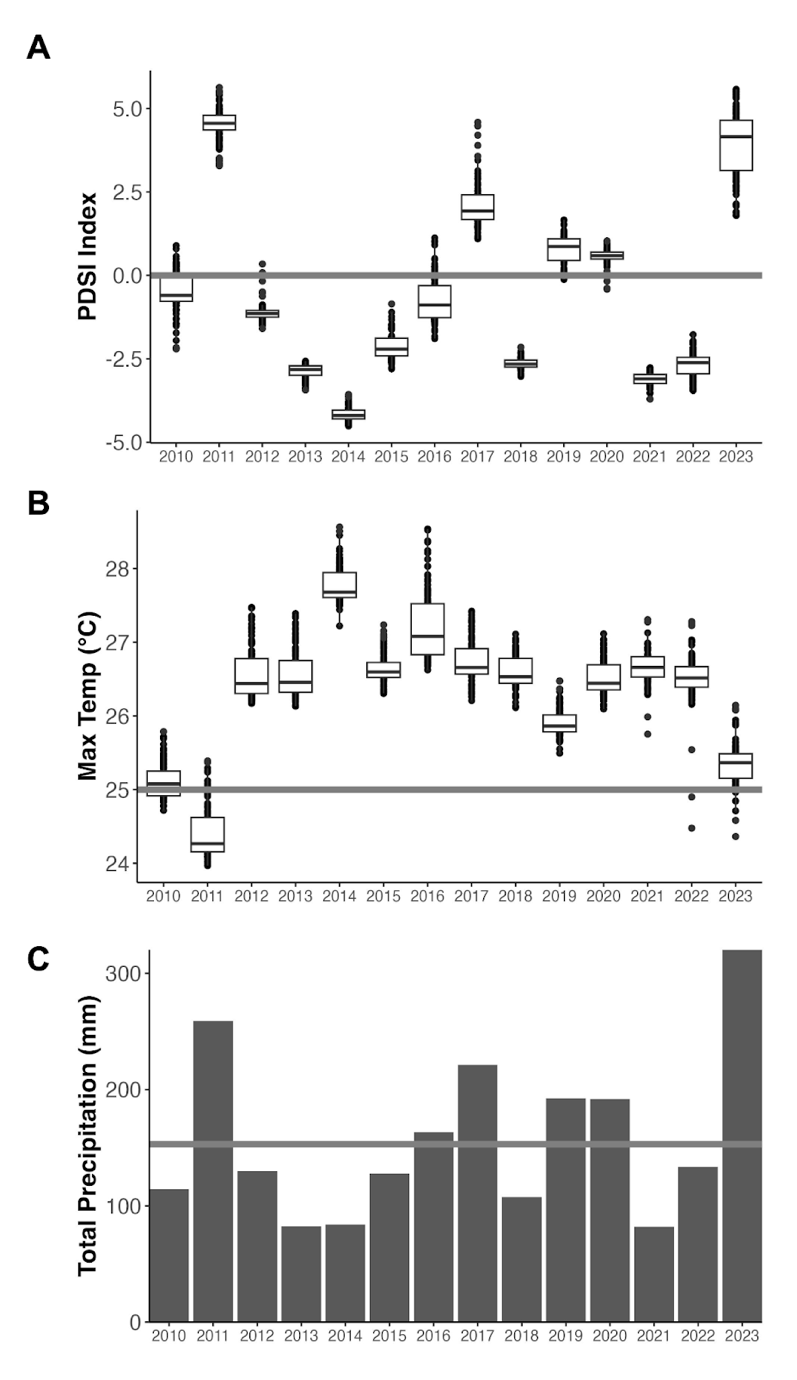

**Figure S2**. **Study site trap station and clusterID information.** Additional information on trap stations (A) some assigned clusterIDs, (B) number of unique trap stations per clusterID, and (C) associated land cover type. Land cover data comes from California's Department of Conservation Farmland Mapping and Monitoring Program (FMMP; https://www.conservation.ca.gov/dlrp/fmmp). In Figure 2C, each trap point is defined by the land cover type beneath it. These include natural, peri-urban, primary agriculture, secondary agriculture, and urban land cover types.

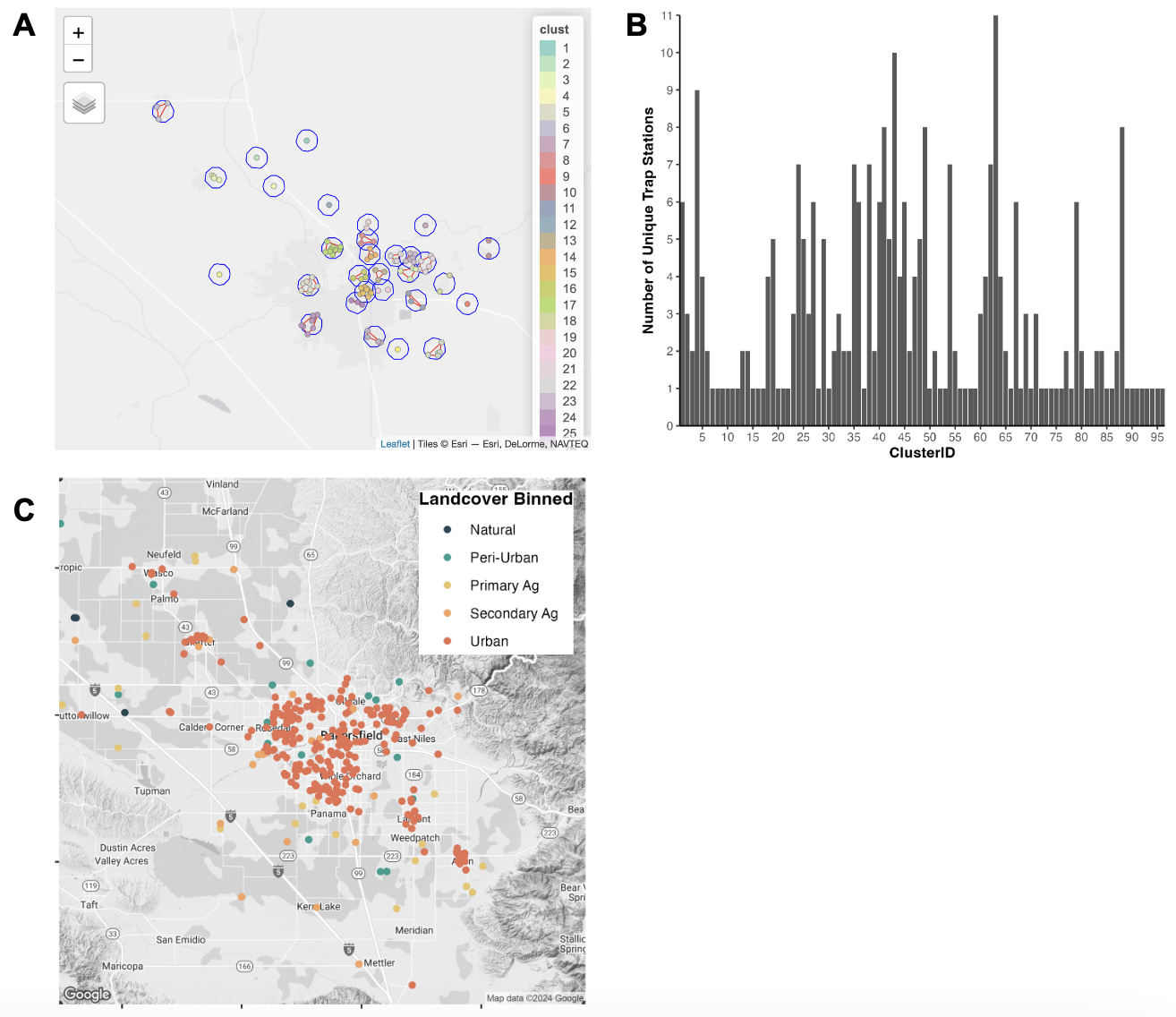

**Figure S3**. **Pesticide application in Kern County.** Public health pesticide application of the log_10_ transformed (A) larvicide and (B) adulticide in pounds per month-year of the entire county.

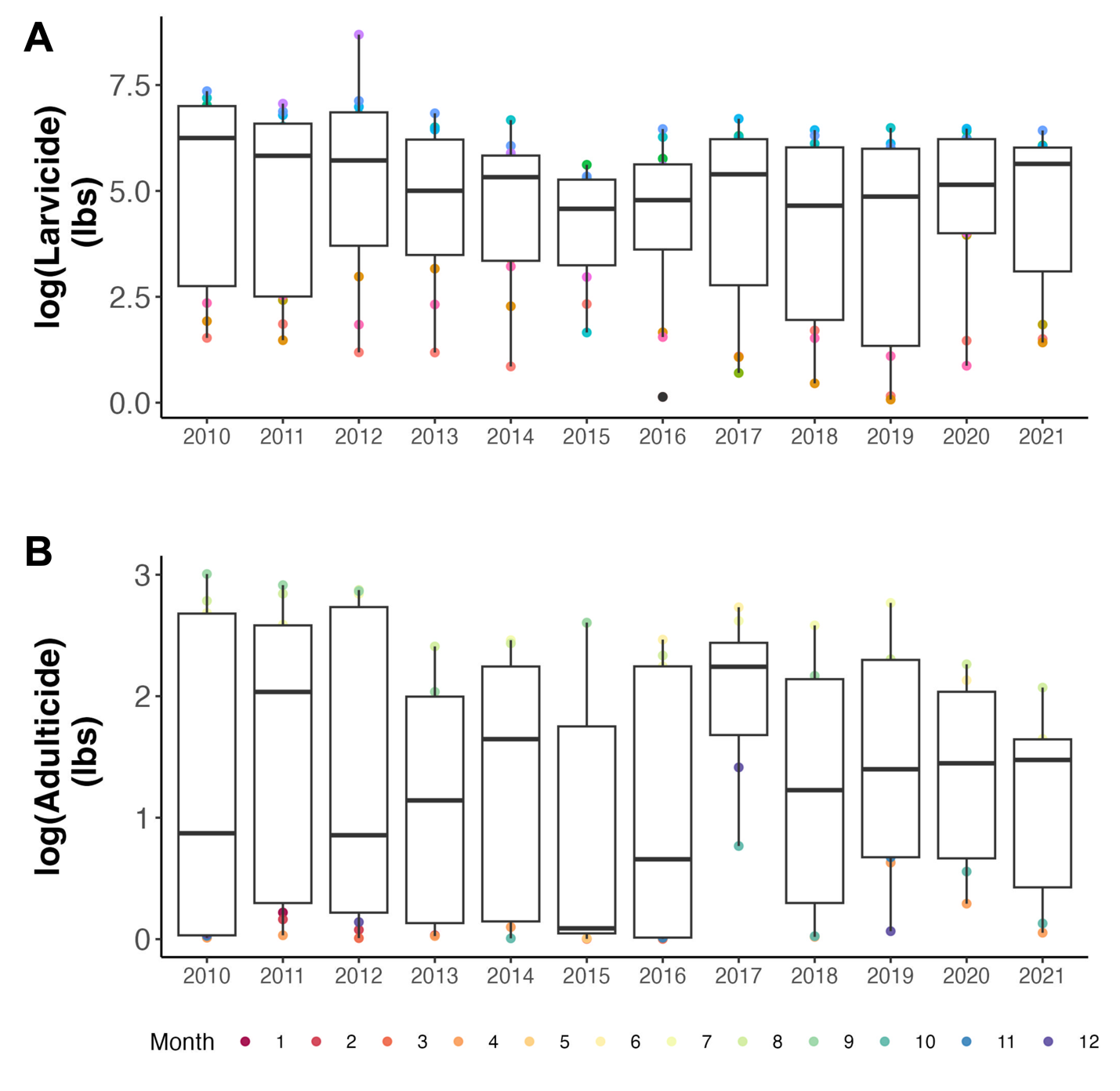

**Figure S4**. **Coefficient estimates for abundance models with different fixed effects.** Coefficient estimates and 95% confidence intervals for monthly mosquito **abundance models** with **different fixed effects**. Panels are split by mosquito species (top – *Cx. tarsalis*, bottom – *Cx. quinquefasciatus*). Colors correspond to an individual model (covariates and fixed effect terms).

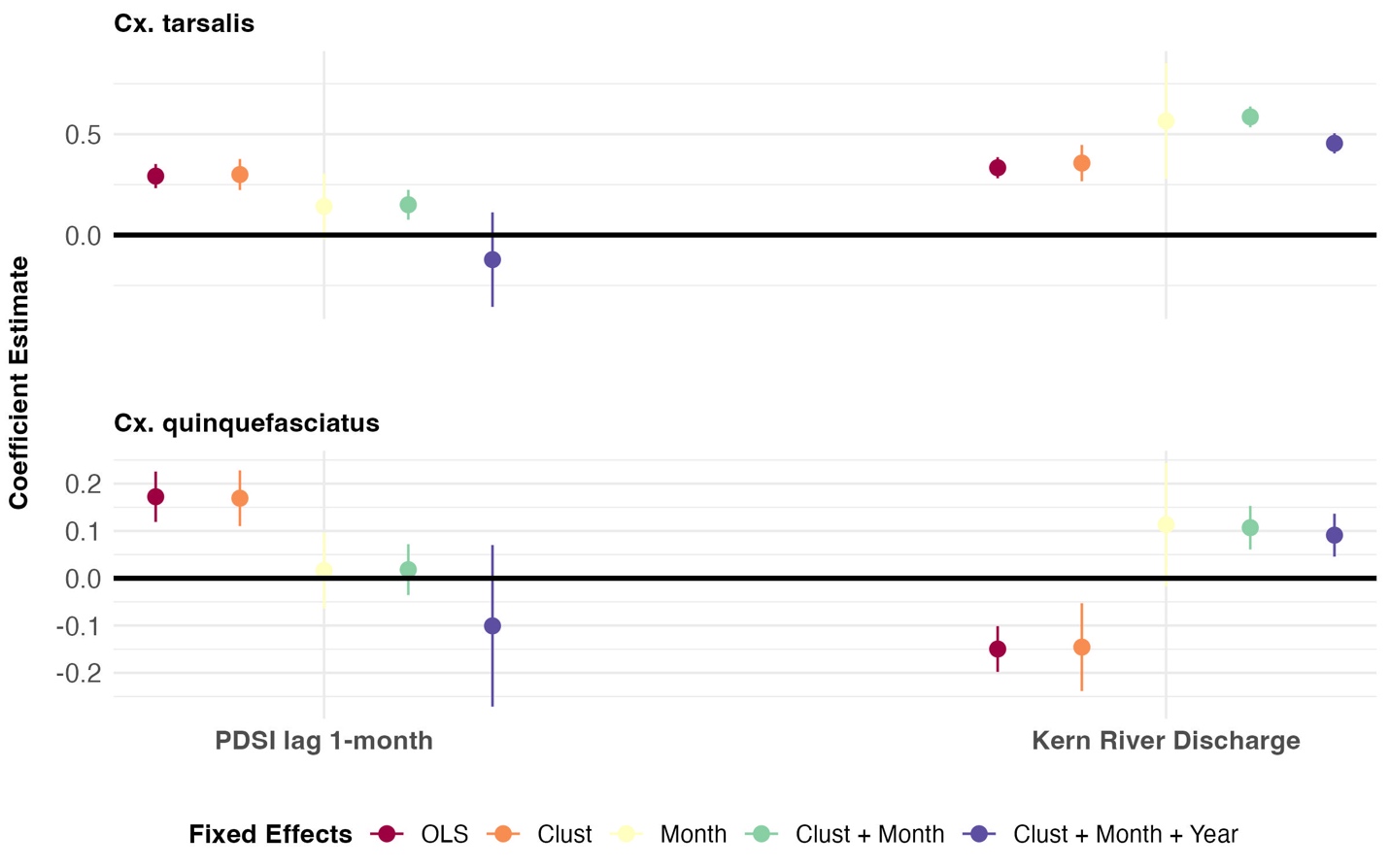

**Figure S5**. **Coefficient estimates for WNV MIR models with different fixed effects.** Coefficient estimates and 95% confidence intervals for monthly mosquito **WNV minimum infection rate (MIR) models** with **different fixed effects**. Panels are split by mosquito species (top – *Cx. tarsalis*, bottom – *Cx. quinquefasciatus*). Colors correspond to an individual model (covariates and fixed effect terms).

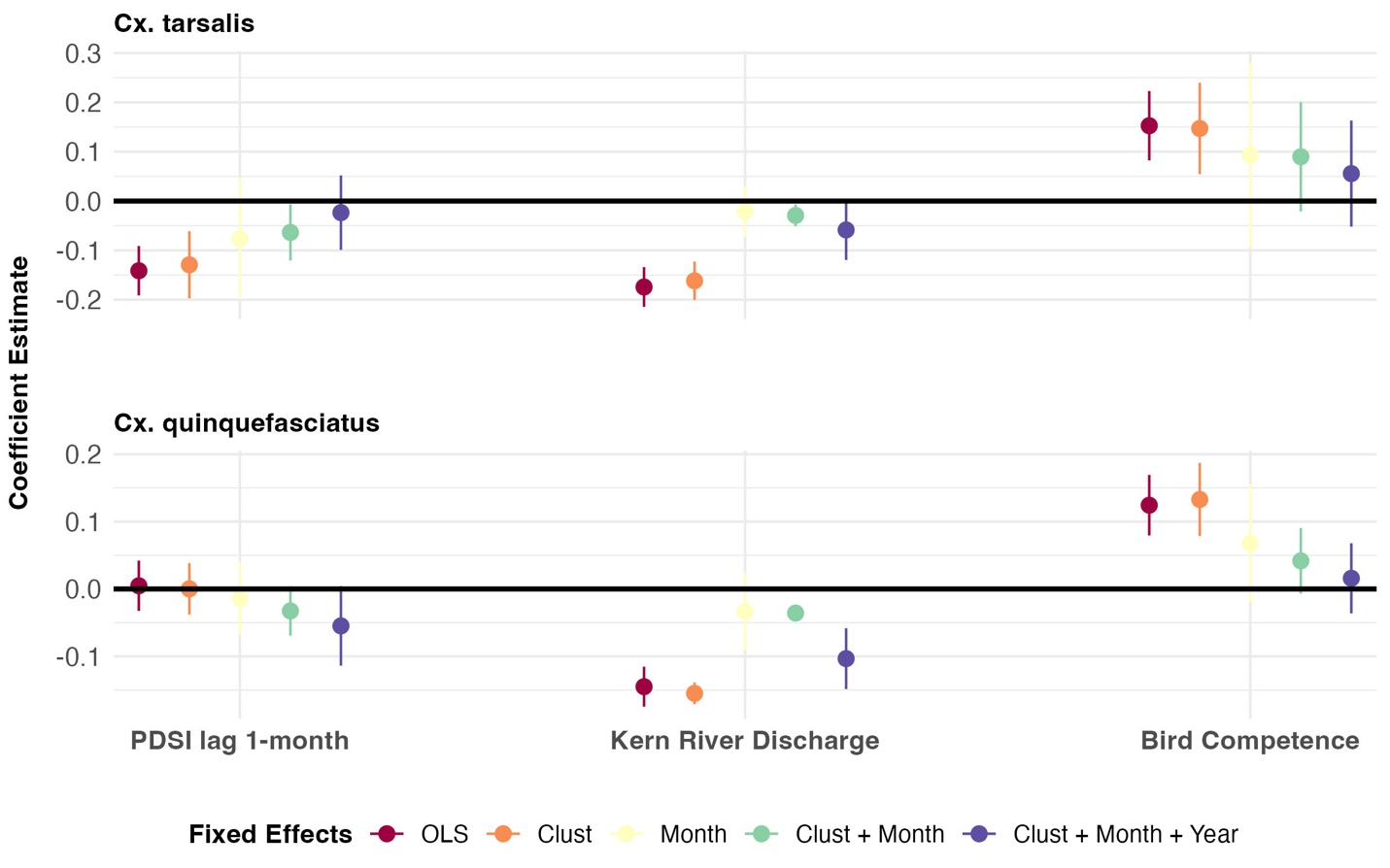

**Figure S6.** **Coefficient estimates for abundance models with different Conley standard errors.** Coefficient estimates and 95% confidence intervals for monthly mosquito **abundance models** with the fixed effects ‘clustID’ and ‘month’ and varying **Conley standard errors**. Panels are split by mosquito species (top – *Cx. tarsalis*, bottom – *Cx. quinquefasciatus*). Colors correspond to an individual model (covariates and varying Conley standard errors). For the final model a Conley standard error of 11 km was used.

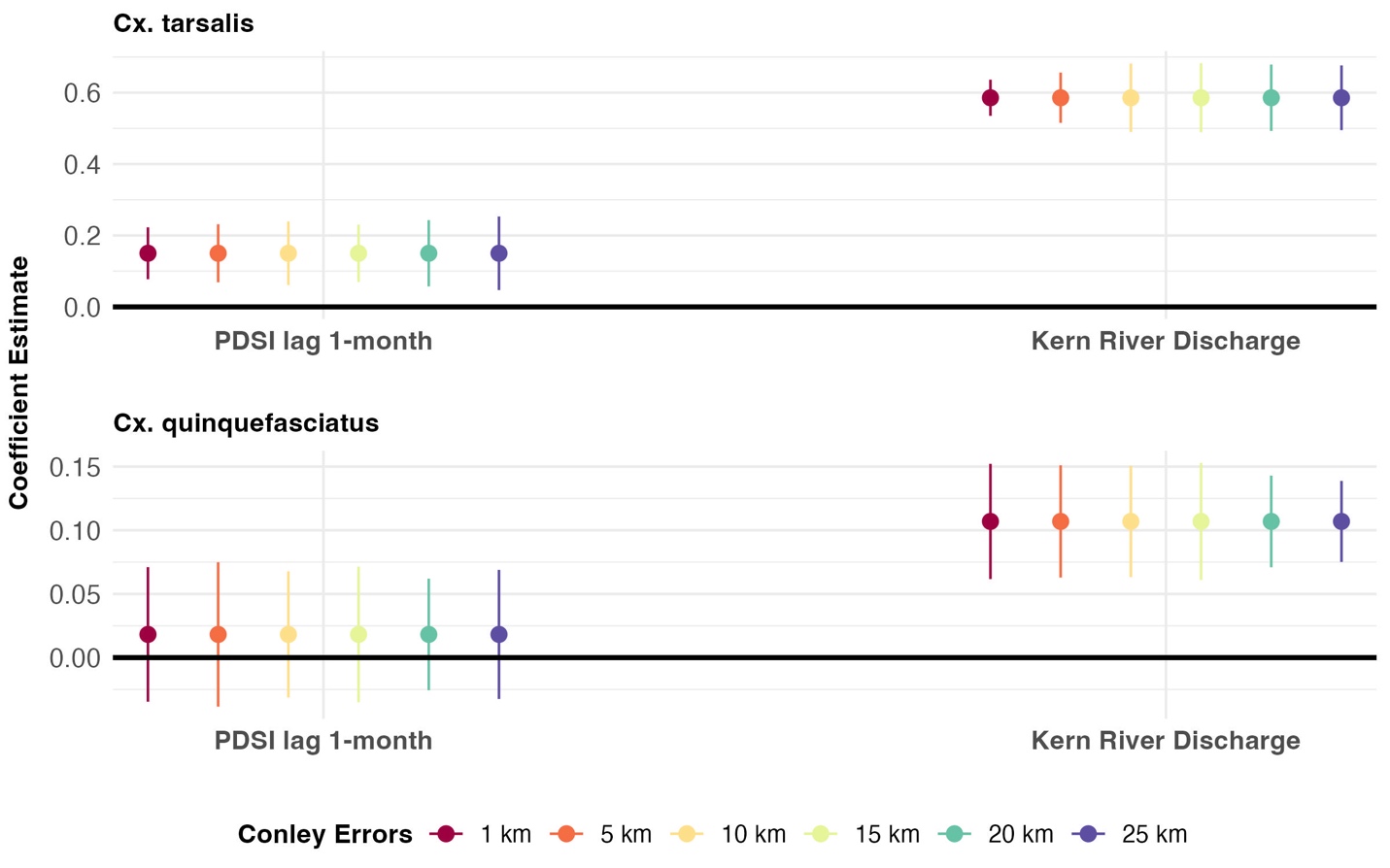

**Figure S7. Coefficient estimates for WNV MIR models with different Conley standard errors.** Coefficient estimates and 95% confidence intervals for monthly mosquito **WNV minimum infection rate (MIR) models** with the fixed effects ‘clustID’ and ‘month’ and varying **Conley standard errors**. Panels are split by mosquito species (top – *Cx. tarsalis*, bottom – *Cx. quinquefasciatus*). Colors correspond to an individual model (covariates and varying Conley standard errors). For the final model a Conley standard error of 11 km was used.

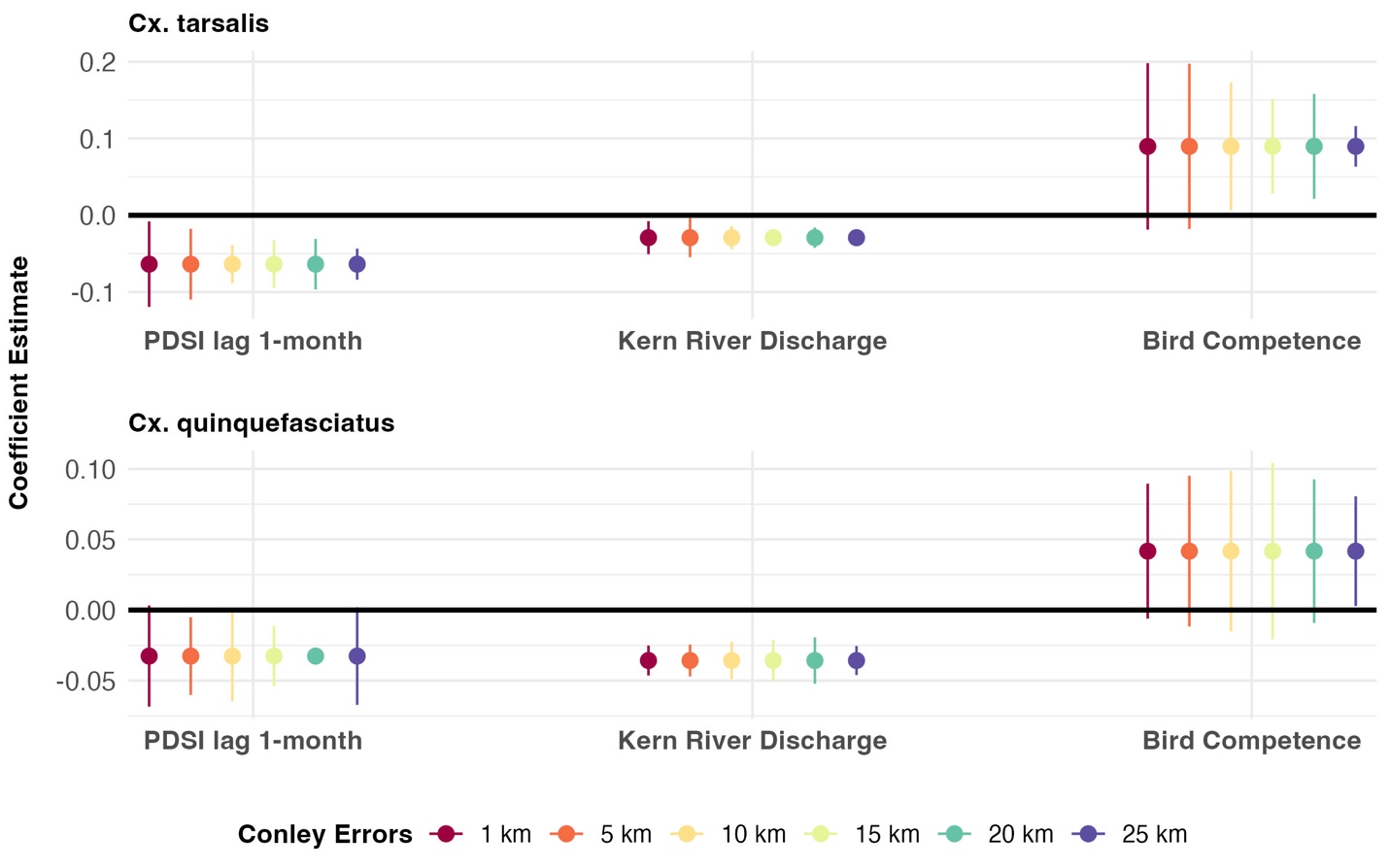

**Figure S8. Coefficient estimates for abundance models with different types of standard errors.** Coefficient estimates and 95% confidence intervals for monthly mosquito **abundance models** with the fixed effects ‘clustID’ and ‘month’ and varying **standard error calculations**. Panels are split by mosquito species (top – *Cx. tarsalis*, bottom – *Cx. quinquefasciatus*). Colors correspond to an individual model with different standard errors calculated in the `sandwich` package (Zeileis et al. 2006).

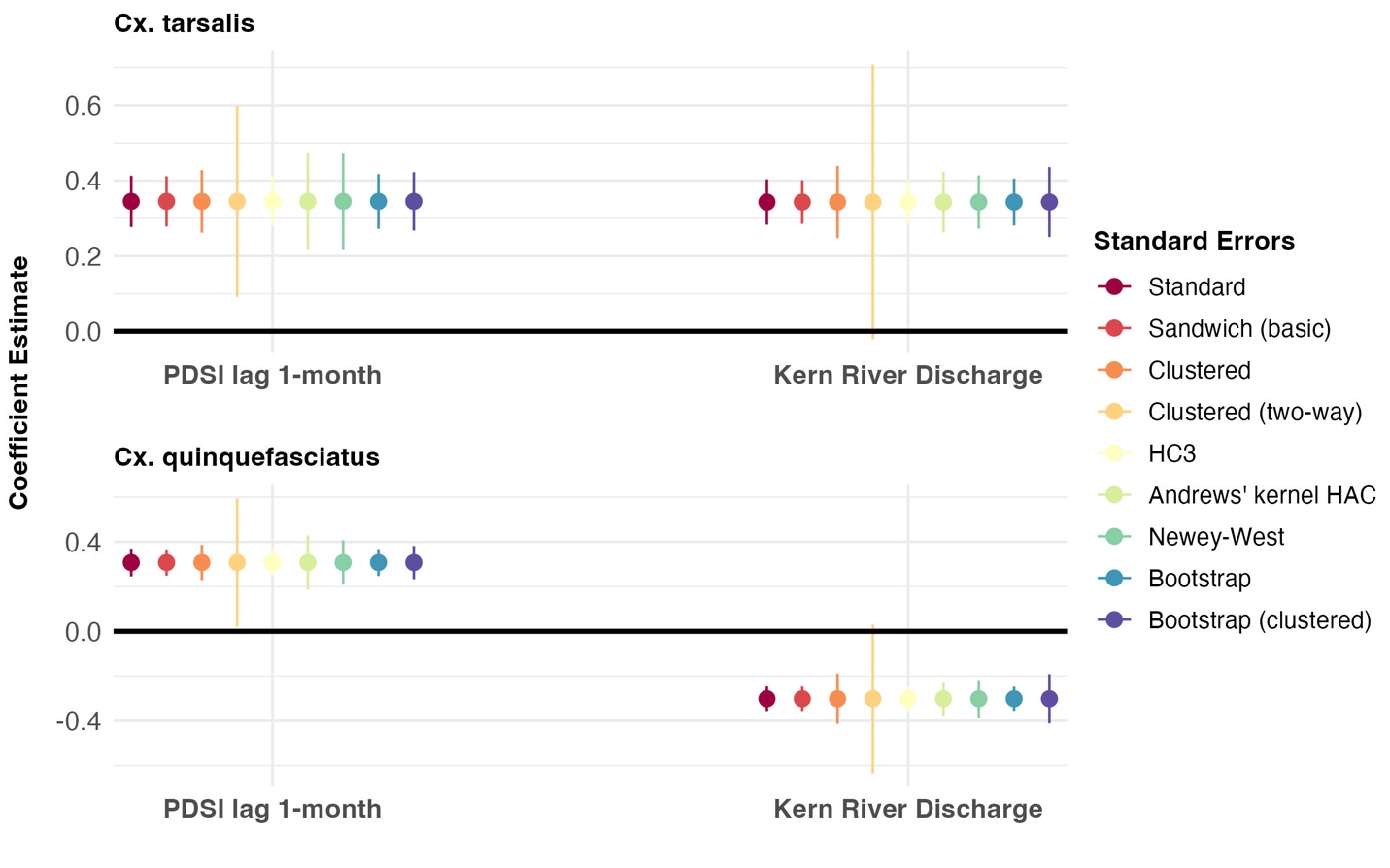

**Figure S9. Coefficient estimates for WNV MIR models with different types of standard errors.** Coefficient estimates and 95% confidence intervals for monthly mosquito **WNV minimum infection rate (MIR) models** with the fixed effects ‘clustID’ and ‘month’ and varying **standard error calculations**. Panels are split by mosquito species (top – *Cx. tarsalis*, bottom – *Cx. quinquefasciatus*). Colors correspond to an individual model with different standard errors calculated in the `sandwich` package (Zeileis et al. 2006).

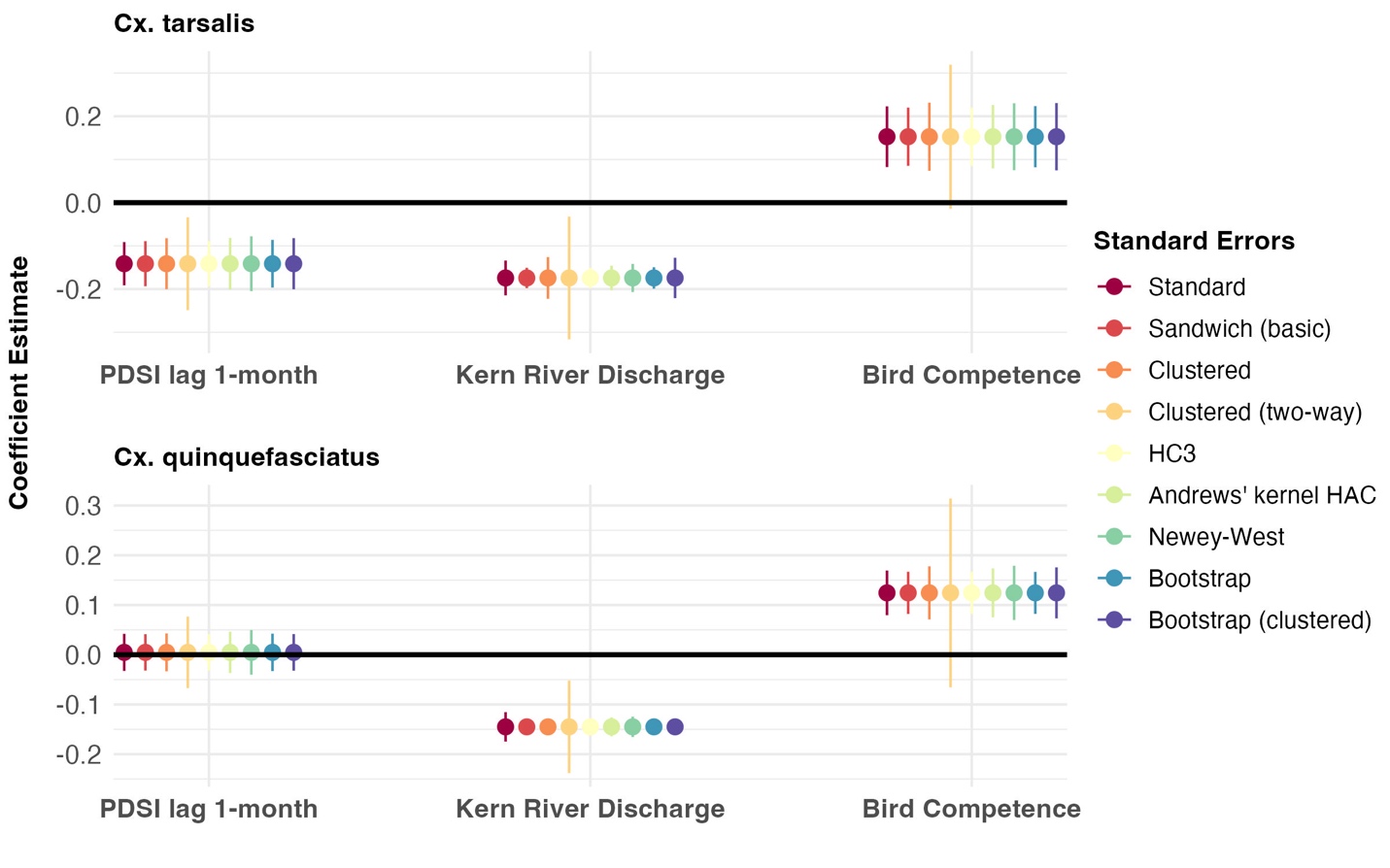
